# Both default and multiple-demand regions represent semantic goal information

**DOI:** 10.1101/2020.07.09.196048

**Authors:** Xiuyi Wang, Zhiyao Gao, Jonathan Smallwood, Elizabeth Jefferies

**Affiliations:** Department of Psychology, University of York, Heslington, York YO10 5DD, United Kingdom

**Keywords:** Semantic, control, default mode network, multiple demand, feature representation, decoding

## Abstract

While the multiple-demand network plays an established role in cognitive flexibility, the role of default mode network is more poorly understood. In this study, we used a semantic feature matching task combined with multivoxel pattern decoding to test contrasting functional accounts. By one view, default mode and multiple-demand networks have opposing roles in cognition; consequently, while multiple-demand regions can decode current goal information, semantically-relevant default network regions might decode conceptual similarity irrespective of task demands. Alternatively, default mode regions might show sensitivity to changing task demands like multiple-demand regions, consistent with evidence that both networks dynamically alter their patterns of connectivity depending on the context. Our task required participants to integrate conceptual knowledge with changing task goals, such that successive decisions were based on different features of the items (colour, shape and size). This allowed us to simultaneously decode semantic category and current goal information using a whole-brain searchlight decoding approach. As expected, multiple-demand regions represented information about the currently-relevant conceptual feature, yet similar decoding results were found in default mode network regions, including angular gyrus and posterior cingulate cortex. Semantic category irrespective of task demands could be decoded in lateral occipital cortex, but not in most regions of default mode network. These results show that conceptual information related to the current goal dominates the multivariate response within default mode network. In this way, default mode network nodes support flexible memory retrieval by modulating their response to suit active task goals, alongside regions of multiple-demand cortex.

**Significance Statement:** We tested contrasting accounts of default mode network (DMN) function using multivoxel pattern analysis. By one view, semantically-relevant parts of DMN represent conceptual similarity, irrespective of task context. By an alternative view, DMN tracks changing task demands. Our semantic feature matching task required participants to integrate conceptual knowledge with task goals, such that successive decisions were based on different features of the items. We demonstrate that DMN regions can decode current goal, alongside multiple-demand regions traditionally associated with cognitive control. The successful decoding of goal information plus largely absent category decoding effects within DMN indicates that this network supports flexible semantic cognition.

## 1. Introduction

Human cognition is flexible, enabling us to select appropriate information from memory, according to our current goals. Accounts of cognitive flexibility emphasise the role of the multiple demand network (MDN; Duncan, 2010), which shows stronger responses in demanding conditions across tasks (Fedorenko et al., 2013; Turnbull et al., 2019) and activation patterns that can classify different task-critical details in an adaptive fashion (Erez and Duncan, 2015; Cole et al., 2016; Bracci et al., 2017; Qiao et al., 2017). Yet the role of MDN in cognitive flexibility is typically assessed using a regions-of-interest (ROIs) approach (Waskom et al., 2014; Erez and Duncan, 2015; Cole et al., 2016; Bracci et al., 2017; Qiao et al., 2017); consequently, the contribution of other heteromodal brain regions remains unclear.

A key debate concerns the role of default mode network (DMN) in cognition. DMN and MDN typically show opposing responses to task difficulty and negative functional connectivity at rest (Mckiernan et al., 2003; Fox et al., 2005; Kelly et al., 2008; Fornito et al., 2012). Nevertheless, semantically-relevant DMN regions, including left angular gyrus and lateral temporal cortex, show less deactivation, relative to rest, when semantic and non-semantic tasks are compared (Binder et al., 1999; 2009; Humphreys et al., 2015) – even when task difficulty is taken into account (Wirth et al., 2011; Binder et al., 2005; Murphy et al., 2018; Seghier et al., 2010). DMN is highly heteromodal, and thought to support information integration (Simony et al., 2016; Lanzoni et al., 2020), relevant to episodic retrieval (Sestieri et al., 2011) and semantic cognition (Binder & Desai, 2011; Wirth et al., 2011). These observations suggest that semantic DMN regions might extract information about global conceptual similarity in long-term memory (Murphy et al., 2017; Wang et al., 2020). By this view, patterns of response within DMN regions might be driven by conceptual similarity, irrespective of task.

By an alternative view, DMN might be sensitive to changing task demands and support cognition in a more active and flexible way. DMN is situated at the top of a cortical hierarchy, exhibiting the greatest distance (both physically and in connectivity space) from primary sensory/motor regions (Margulies et al., 2016). On this heteromodal to unimodal gradient, frontoparietal control and DMN regions occupy adjacent locations and have similar representational content (González-García et al., 2018). DMN has broad patterns of connectivity (Braga et al., 2013), which are dynamically altered depending on fluctuations in task demands (Vatansever et al., 2015, 2017) even as this network deactivates (Krieger-Redwood et al., 2016). These functional interactions positively correlate with behavioural performance (Krieger-Redwood et al., 2015; Vatansever et al., 2017). Moreover, DMN supports detailed thoughts about demanding tasks as well as off-task states (Sormaz et al., 2018). This network is engaged when participants are preparing for a task (Crittenden et al., 2015; Smith et al., 2018) or applying task rules from memory (Vatansever et al., 2017). Collectively, these observations suggest DMN might maintain currently-relevant information that can bias ongoing processing, rather than always reflecting conceptual similarity in long-term memory (Crittenden et al., 2015; Murphy et al., 2017; Smith et al., 2018).

In the present study, we contrasted these accounts to establish the contribution of DMN to controlled semantic cognition. We examined whether patterns of response across DMN capture long-term semantic similarity or the short-term behavioural relevance of specific semantic features. Participants were asked to match items from three categories (animals, plants, tools) according to colour, shape or size, with the goal switching on each trial. This allowed us to directly contrast decoding of long-term semantic similarity and goal-relevant features. We conducted whole brain classification analysis to look at task engagement and related the decoding results to localizer tasks used to define DMN and MDN. To anticipate, our results indicate that goal information can be decoded in both MDN and DMN, while lateral occipital cortex is the only region that represents both goal and category information.

## 2. Methods

### 2.1. Participants

The research was approved by the York Neuroimaging Centre and Department of Psychology ethics committees. 31 healthy adults were recruited from the University of York (26 females; age: mean ± SD = 20.60 ± 1.68, range: 18 – 25 years). All participants were right-handed, native English speakers, with normal or corrected-to-normal vision and no history of psychiatric or neurological illness. All volunteers provided written informed consent. One participant with incomplete data (only one of two sessions) was removed. Two more participants were removed because of low accuracy (3SD below the mean). This study provides new analyses of a dataset first reported by Wang et al. (2020).

### 2.2. Design and procedure

Participants completed two fMRI sessions: in the first session, they performed a semantic feature matching task, in which we varied the goal (i.e. the feature to be matched) and the semantic category of the probe word. This allowed us to directly contrast the decoding of goal-relevant features and long-term semantic similarity. In the second session, participants completed easy and hard spatial working memory and arithmetic tasks (from Blank, Kanwisher, & Fedorenko, 2014; Fedorenko et al., 2011, 2013) designed to localise MDN and DMN. The contrast of hard versus easy versions of these tasks robustly activates MDN regions (Fedorenko et al., 2013; Blank et al., 2014), while the easy versus hard contrast activates DMN (Fedorenko et al., 2013; Leech et al., 2011; Mckiernan et al., 2003). In this way, we could establish the overlap between regions representing goal and category information and the MDN and DMN.

### 2.3. Behavioural tasks

#### 2.3.1. Semantic feature matching task

Participants matched probe and target concepts (presented as words) according to a particular semantic feature (colour, shape or size), specified at the start of each trial in a rapid event-related design (Figure 1). Two third of trials were matching trials in which probe and target shared the target feature (i.e. Colour: strawberry and cherry are both red) and one third were non-matching trials in which probe and target do not share similar target feature (i.e. Colour: lemon and raspberry have different colours, although they are semantically related). Participants pressed a button with their right index finger to indicate a matching trial and responded with their right middle finger to indicate a non-matching trial. All the probe words belonged to one of three categories: animal, tool, and plant. This gave rise to 9 combinations of goal feature and probe category, with 36 items for each condition. These trials were divided evenly into 4 runs. The order of runs and trials within each run was randomized across subjects. Each run lasted for 600s. The target words were drawn from a wider range of categories than the probe words. Probe and target words were matched on word length (number of letters), word frequency (based on SUBTLEX-UK: Subtitle-based word frequencies for British English) (van Heuven et al., 2014) and word concreteness (Brysbaert, Warriner, & Kuperman, 2014) across conditions, respectively. Participants were provided with feedback during task training but not during the main experiment.

**Figure 1:**
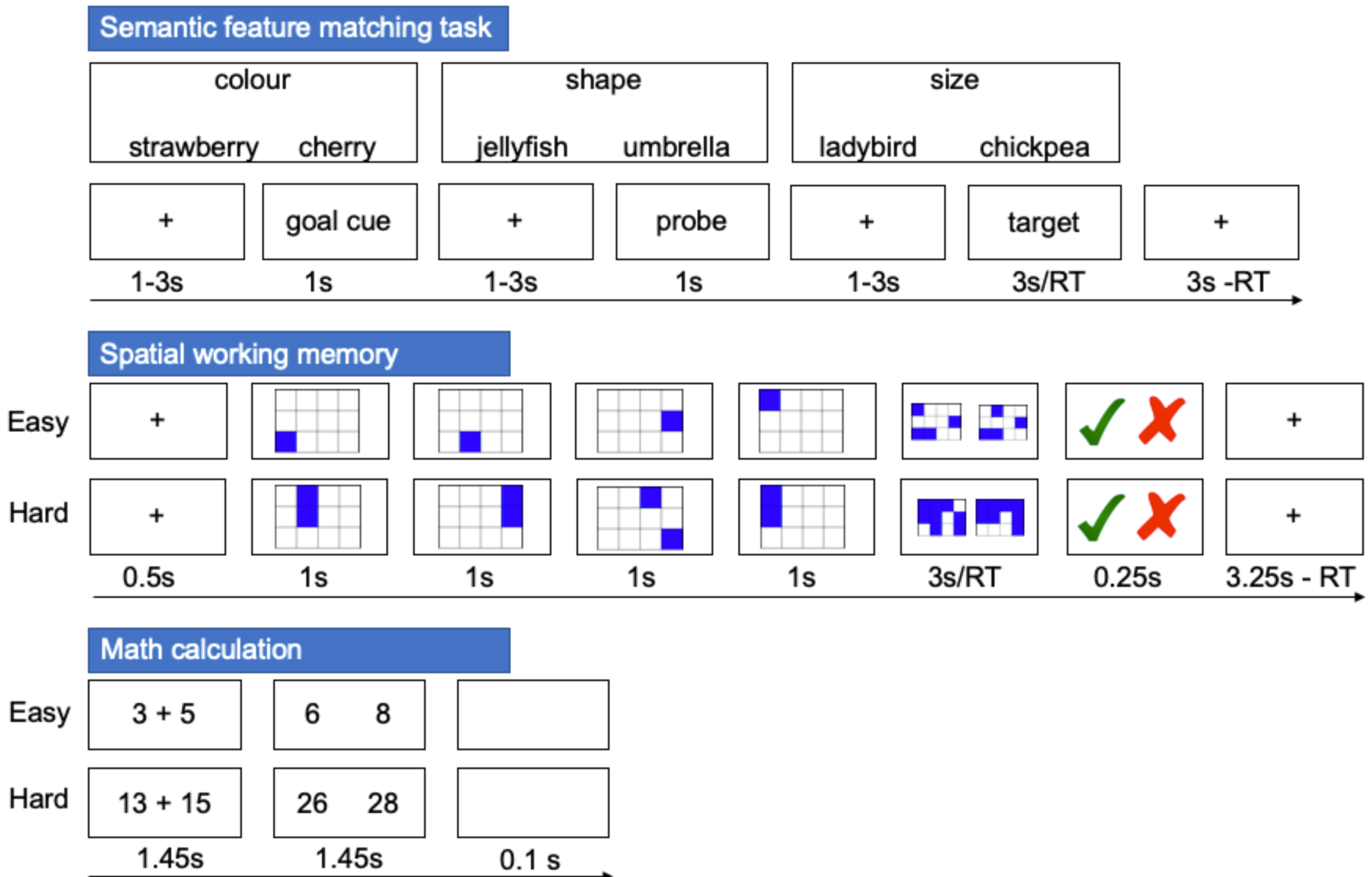
Illustration of semantic feature matching task, spatial working memory task and math task.

In order to maximize the statistical power of the rapid event-related fMRI data analysis, the stimuli were presented with a temporal jitter, randomized from trial to trial (Dale, 1999). The inter-stimulus intervals (between goal cue and probe word, and probe and target word) and the inter-trial interval varied from 1 to 3s. Each trial started with a fixation, followed by a goal cue slide specifying the feature to match (colour, shape or size), presented for 1s. This was followed by the second fixation and then the probe word, presented for 1s. Finally, there was the third fixation followed by the target word, triggering the onset of the decision-making period. The target remained visible until the participant responded, or for a maximum of 3s. The goal cue, probe, and target words were presented centrally on the screen. Both response time and accuracy were recorded.

A trial consisted of three events: (1) A goal cue which indicated the relevant feature for the trial, (2) a probe word, and (3) a target word, which was followed by a response. Based on these events, we separated each trial into three different time periods: a ‘goal cue period’, a ‘probe word period’, and a ‘target word period’. Our main analysis concerned the probe word period since, for this time point, we were able to perform both goal and category decoding for the same items. We also performed secondary analyses using the goal cue period and the target word period (the category classifier could not be applied at these time points; see below).

#### 2.3.2. Spatial working memory task

Participants had to keep track of four or eight sequentially presented locations in a 3×4 grid (Fedorenko et al., 2011), giving rise to easy and hard spatial working memory conditions. Stimuli were presented at the center of the screen across four steps. Each of these steps lasted for 1s and highlighted one location on the grid in the easy condition and two locations in the hard condition. This was followed by a decision phase, which showed two grids side by side. One grid contained the locations shown on the previous four steps, while the other contained one or two locations in the wrong place. Participants indicated their recognition of these locations in a two-alternative, forced-choice paradigm via a button press and feedback was immediately provided. Each run consisted of 12 experimental blocks (6 blocks per condition and 4 trials in a 32 s block) and 4 fixation blocks (each 16s long), resulting in a total time of 448s. This task included two runs containing the two conditions, in a standard block design. Condition order was counterbalanced across runs, and run order was counterbalanced across participants for each task.

#### 2.3.3. Math task

In this task, participants performed addition with smaller or larger numbers, giving rise to easy and hard conditions. Participants saw an arithmetic expression on the screen for 1.45s and were then given two numbers as potential answers, for 1.45s. Each trial ended with a blank screen lasting for 0.1s. Each run consisted of 12 experimental blocks (with 4 trials per block) and 4 fixation blocks, resulting in a total time of 316s. This task included two runs containing the two conditions, presented in a standard block design. Condition order was counterbalanced across runs and run order was counterbalanced across participants for each task. All the stimuli were presented using Psychopy (Peirce, 2007).

### 2.4. fMRI data acquisition

Structural and functional data were collected on a Siemens Prisma 3T MRI scanner at the York Neuroimaging Centre. The scanning protocols included a T1-weighted MPRAGE sequence with whole-brain coverage. The structural scan used: acquisition matrix of 176 × 256 × 256 and voxel size 1 × 1 × 1mm^3^, repetition time (TR) = 2300ms, and echo time (TE) = 2.26ms. Functional data were acquired using an EPI sequence with an 80^0^ flip angle and using GRAPPA with an acceleration factor of 2 in 3 x 3 x 4mm voxels in 64-axial slices. The functional scan used: 55 3-mm-thick slices acquired in an interleaved order (with 33% distance factor), TR = 3000ms, TE = 15ms, FoV = 192mm.

### 2.5. MRI data pre-processing

Functional and structural data pre-processing for classification was carried out using FMRIB’s Software Library (FSL version 6, fsl.fmrib.ox.ac.uk/fsl/fslwiki/FEAT/). The T1-weighted structural brain images were extracted. Structural images were registered to the MNI-152 template using FMRIB’s linear image registration tool (FLIRT). fMRI data pre-processing included motion correction, slice-timing correction, and high-pass filtering at 100s. Motion-affected volumes were detected and then were fully removed from the fMRI data (using scrubbing; Power, Barnes, Snyder, Schlaggar, & Petersen, 2012). No spatial smoothing was applied at this point to preserve fine-grained patterns of voxel activations (Haynes and Rees, 2006).

### 2.6. MRI data analysis

Classification analysis focussed on the probe word period when both goal and semantic category were manipulated. First, a univariate analysis was used to identify the regions showing stronger activation when performing the task relative to rest. Secondly, we conducted a univariate analysis to establish the parameter estimates for each feature and each category. Next, whole-brain searchlight analysis revealed regions in which the multivariate response classified the goal and/or semantic category. We also performed supplementary whole-brain searchlight decoding analyses during the goal cue and target word periods. These analyses examined goal but not category decoding since the goal cues preceded the presentation of concepts, and category was not manipulated for the target word.

#### 2.6.1. Univariate analysis of the semantic feature matching task

To estimate the effects of task demands during the semantic feature matching task, we built a general linear model. Spatial smoothing with a 5mm FWHM Gaussian filter was applied. The main effect of task was modelled as epochs lasting from the target onset to response, thus controlling for lengthened BOLD responses on trials with longer response times. We also modelled the instruction period, two inter-stimulus interval periods, probe word period, and incorrect trials as regressors of no interest. We included the within-trial inter-stimulus intervals as regressors of no interest (removing them from the implicit baseline) since participants were maintaining feature and probe information during these two periods. Six head motion parameters were further included in the GLM as regressors of no interest to control for potential confounding effects of head motion. The four runs of each task were included in a fixed-effects model for each individual. Group-level analyses were carried out using a cluster-forming threshold of Z > 3.1, with p < 0.05 (corrected for family-wise error (FWE) rate using random field theory).

#### 2.6.2. First-level univariate model of probe word period for classification analysis

Univariate models were used to extract the β value for each feature (colour, shape and size), for each run, in each voxel of the brain for goal decoding. The resulting pre-processed unsmoothed data were analysed using first-level FEAT (v6.00; fsl.fmrib.ox.ac.uk/fsl/fslwiki) in subject native space. A general linear model was built for each run for each participant. One regressor was created for each feature. Trials were modelled as epochs lasting from probe onset to probe offset convolved with a canonical hemodynamic response function (HRF). We modelled other time points as regressors of no interest, including the goal cue period from goal cue word onset to offset, the target word period from target word onset to response, the two inter-stimulus interval periods and the incorrect trials. We also ran the GLM without these two inter-stimulus intervals and the results were largely unchanged. We included temporal autocorrelation correction and six motion parameters as nuisance covariates. Within each run, time points were averaged across trials within each feature, such that there were four data points per feature per subject (one per run) (for similar implementations, see Cole, Ito, & Braver, 2016; Connolly et al., 2012). To increase the statistical power, we also created more data points by randomly but evenly dividing all the correct trials of each feature in each run into three sub-datasets (for similar implementations, see Hebart, Bankson, Harel, Baker, & Cichy, 2018). This resulted in three data points per feature per run per subject (e.g. nine observations), which were z-scored within each run at each voxel (Hanke et al., 2009; Connolly et al., 2012).

Univariate models were also used to extract the β value for each category (animal, tool and plant), for each run, in each voxel of the brain for category decoding. The procedure for extracting the beta weights for each category reproduced the methods above, except that one regressor was created for each category, and within each run, time points were averaged across trials within each category, such that there were four data points per category per subject (one per run) (for similar implementations, Cole, Ito, & Braver, 2016; Connolly et al., 2012). To increase the statistical power, we also created more data points by randomly but evenly dividing all the correct trials of each category of each run into three sub-datasets (for similar implementations, see Hebart et al., 2018). This resulted in three data points per feature per run per subject (e.g. nine observations), which were z-scored within each run for each voxel.

#### 2.6.3. Decoding analysis of goal information during probe word period

We performed a whole-brain searchlight decoding analysis to investigate which regions represent goal feature information (i.e. colour; shape; size). Classifiers were trained and tested on individual subject data transformed into MNI standard space using the z-scored β-values for each feature. Classification training and testing were done using a leave-one-run-out cross-validation strategy. There were twenty-seven total brain patterns (nine per feature) to train the classifier and nine total brain patterns (three per feature) to test the classifier each time. We tested the discriminability of patterns for the three goals using a whole-brain searchlight with a radius of 6mm (number of voxels = 123) (Kriegeskorte et al., 2006) with linear support vector machines (SVMs) (Vapnik and Chapelle, 2000) (LIBSVM, http://www.csie.ntu.edu.tw/~cjlin/libsvm/) implemented within PyMVPA (Hanke et al., 2009). Classification accuracy for each sphere was assigned to the sphere’s central voxel, in order to produce decoding accuracy maps. The resulting accuracy maps were then smoothed with a Gaussian kernel (6mm FWHM). We performed a group analysis as a second-level analysis based on pattern-information maps to determine whether accuracy maps were above chance-levels (accuracy > 0.33) (Kriegeskorte et al., 2006). Individual decoding accuracy maps from all participants were submitted to a nonparametric one-sample t-test on each voxel based on permutation methods implemented using the Randomize tool in FSL (5000 permutations) (https://fsl.fmrib.ox.ac.uk/fsl/fslwiki/Randomise/). Threshold-free cluster enhancement (TFCE; Smith & Nichols, 2009) was used to identify significant clusters from the permutation tests. The final results were thresholded at a TFCE cluster-corrected p-value < 0.05 after controlling for family-wise error rate.

#### 2.6.4. Decoding analysis of category information during probe word period

We performed a second whole-brain searchlight decoding analysis to investigate whether there are any regions that represent category information not required for the task. The procedures were as above, except we extracted the z-scored β-values for each category (animal; tool; plant) for training and testing.

#### 2.6.5. Control feature decoding analysis of goal cue and target word periods

Our main analysis examined the probe word period, which allowed decoding of both goal and semantic category. To understand the nature of goal representation during the probe word period, we also performed secondary analyses of the goal cue and target word periods, examining the decoding of goal information. The goal cue period was modelled using a fixed duration (1s). For the target word period, as there were differences in response time across conditions, we included this variable as a parametric regressor in the GLM for all trials with a fixed duration (1s) (for similar implementations, see Erez & Duncan, 2015; Todd, Nystrom, & Cohen, 2013; Waskom et al., 2014). In addition, we used another method to control for lengthened BOLD response as a result of longer response times. We modelled the event duration using a variable epoch approach, from target onset to response. We then computed the voxelwise beta patterns and did the whole-brain searchlight analysis as before. The results were largely unchanged and are shown in https://osf.io/vn7ws/.

#### 2.6.6. Localizer task analysis

In the spatial working memory task and math tasks, we examined the contrast of hard versus easy to define MDN regions, and the contrast of easy versus hard to define DMN regions. This allowed us to establish whether clusters that could classify category and goal fell within these networks defined using the same participants. The two runs of each task were included in a fixed-effects model for each individual. Group-level analyses were carried out using a cluster-forming threshold of Z > 3.1 and corrected these at p < 0.05 (corrected for FWE rate using random field theory). Results were visualized using BrainNet Viewer (Xia et al., 2013).

To investigate whether regions that represent goal information overlap with MDN or DMN, we compared the goal decoding results with both our localizer contrasts and intrinsic connectivity networks defined by Yeo et al. (2011).

#### 2.6.7. Relationship between behavioural performance and classification results

Since activity in the DMN and MDN is known to be modulated by task difficulty, we tested for the possibility that goal discrimination in these networks is related to behavioural measures of task difficulty. We defined six regions of interest (ROIs) in these two networks by overlapping the goal classification results with the Schaefer parcellation (Schaefer et al., 2018); the ROIs are shown in the Results. We then correlated the classification accuracy in each ROI with the absolute difference in response time between task features. We reasoned that if goal discrimination effects are related to differences in difficulty, we would find significant correlations.

### 2.7. Code accessibility

All summary data, materials and code used in the analysis are accessible in the Open Science Framework at https://osf.io/vn7ws/.

## 3. Results

### 3.1. Behavioural results

There was no significant difference in accuracy across trials probing the three feature goals (F (2, 213) = 1.027, p = 0.360). However, there were significant differences in response time (F(2, 213) = 12.444, p < 0.000). Colour decisions were faster than both shape (p < 0.006) and size decisions (p < 0.0005), which did not differ (p < 0.163). There were no significant differences in accuracy (F (2, 213) = 0.688, p < 0.504) or response times across categories (F(2, 213) = 0.742, p < 0.478) (Figure 2). Figure 2. The behavioural performance of semantic feature matching task (** p < 0.01; *** p < 0.001).

**Figure 2:**
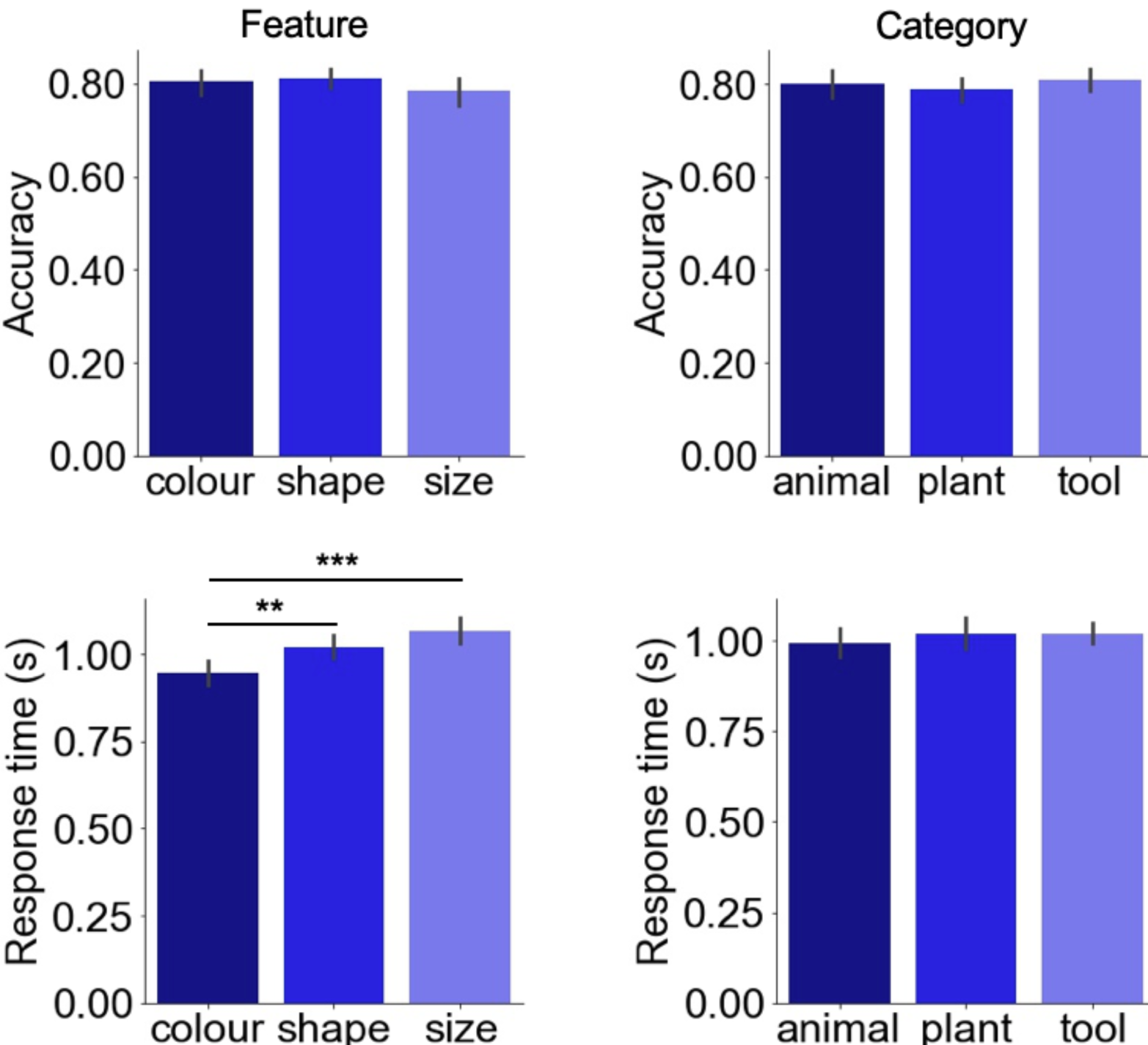
The behavioural performance of semantic feature matching task (**p<0.01;***p<0.001)

### 3.2. Regions involved in semantic feature matching task

We performed a univariate analysis to reveal regions that are involved in the semantic feature matching task. There was activation in MDN and visual regions, including bilateral middle frontal gyrus, bilateral intraparietal sulcus, bilateral pre-supplementary motor area, bilateral medial occipital cortex, left premotor cortex, and left lateral occipital cortex (Figure 3A). The unthresholded map for the semantic feature matching task correlated with the unthresholded MDN localiser maps derived from the spatial working memory task (r = 0.38, p < 0.0001) and math task (r = 0.44, p < 0.0001), demonstrating that semantic feature matching was relatively demanding.

**Figure 3:**
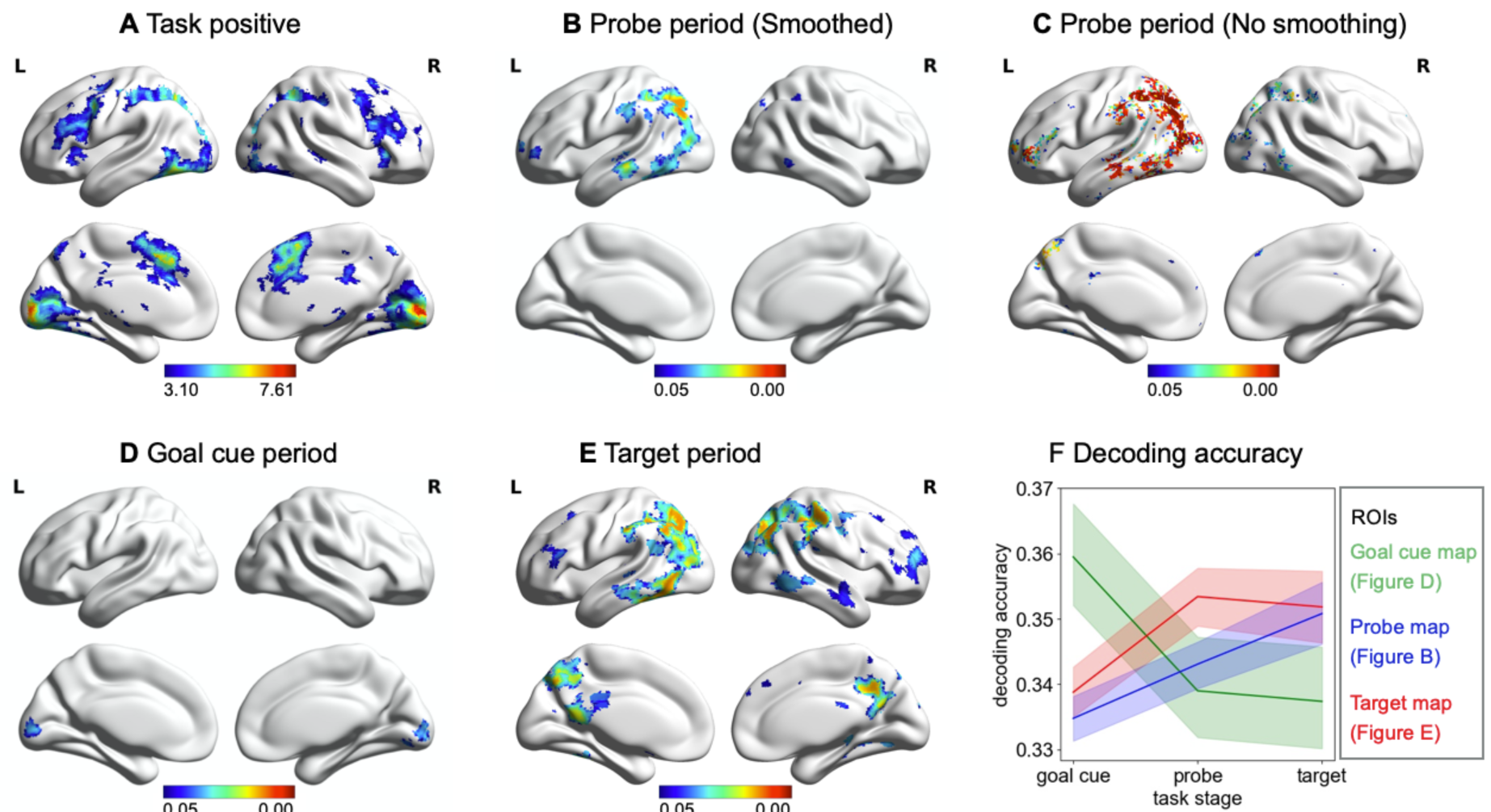
Regions that are involved in the semantic feature matching task and represent goal information for each period. A – Univariate response to the semantic feature matching task (FWE-corrected, p < 0.05). B, C, D, E – Regions that represent goal information for each period (FWE-corrected, p < 0.05). F – Schematic summarising the decoding accuracy of each set of regions found to represent goal information through the task. The green colour represents decoding accuracy within visual regions that decode goal during the cue period. The red colour represents decoding accuracy of regions that can decode goal during the probe period. The blue colour represents decoding accuracy of regions that can decode goal during the target period.

### 3.3. Regions representing goal information during the probe word period

We performed a whole-brain searchlight decoding analysis to identify regions that could represent goal information during the probe word period. We found goal was decoded in left inferior frontal gyrus, left superior parietal lobule, left intraparietal sulcus, left angular gyrus, left lateral occipital cortex, and left posterior middle temporal gyrus (Figure 3B). Given the spatial proximity of DMN and MDN regions, we reran the decoding analysis without smoothing at the group level and found similar results (Figure 3C).

Decoding of goal information during the probe word period might potentially reflect visual or language processes related to the goal cue word (since each goal was defined with a different word). If this were the case, we would expect to see similar goal classification results in the earlier goal cue period, as well as later during the presentation of the probe words, within visual and/or language regions. Alternatively, decoding of goal during the probe word period might reflect rule maintenance and implementation. If this were the case, we reasoned that the classification results would overlap between probe and target periods, when the goal was being maintained and implemented. Therefore, we performed supplementary decoding analyses of the goal cue and target word periods.

In the goal cue period, we found goal information could be decoded in bilateral calcarine sulcus and lingual gyrus (Figure 3D). This likely reflects the early visual representation of the orthographic goal cue words (i.e. the words ‘colour’, ‘shape’ and ‘size’). This decoding map did not overlap with the searchlight analysis of the probe word period, suggesting goal decoding of the probe word period did not reflect specific representations of the goal cue word.

In the target word period, goal information could be decoded in bilateral inferior frontal gyrus, bilateral superior parietal lobe, bilateral intraparietal sulcus, bilateral angular gyrus, bilateral precuneus cortex, bilateral posterior cingulate cortex, bilateral posterior inferior temporal gyrus and bilateral posterior middle temporal gyrus (Figure 3E). There was a 68% overlap in the voxels that could classify goal during the probe and target word periods. Response time was controlled in this analysis by including this variable as a parametric regressor in the GLM for all trials with a fixed duration (1s). Decoding of the target word period yielded similar results when we controlled response time in a different way by modelling the event duration using a variable epoch approach (these results were highly correlated; r = 0.70, p < 0.0001) and were provided at https://osf.io/vn7ws/.

We used spatial correlations to assess the similarity of goal decoding during the goal cue, probe and target word periods. The similarity between the probe and target word periods was higher than between the goal cue and probe word periods. There was a significant correlation between the decoding accuracy maps for the goal cue and probe word period (r = 0.42, p < 0.0001). There was also a correlation between decoding accuracy for probe and target word periods (r = 0.55, p < 0.0001), and this correlation was significantly stronger (z = 39.71, p = 0.0001). To illustrate this pattern, we defined the regions that represent goal information of each period as ROIs and then extracted the decoding accuracy of each ROI in each period (Figure 3F).

### 3.4. Regions classifying goal overlap with both MDN and DMN

To investigate whether regions that represent goal information overlap with MDN or DMN, we compared regions representing goal information to large scale networks defined by a resting-state parcellation of 1000 brains (Yeo et al., 2011). Regions representing goal information during the probe and target periods overlapped primarily with dorsal attention, default mode and frontoparietal and visual networks (Figure 4).

**Figure 4.**
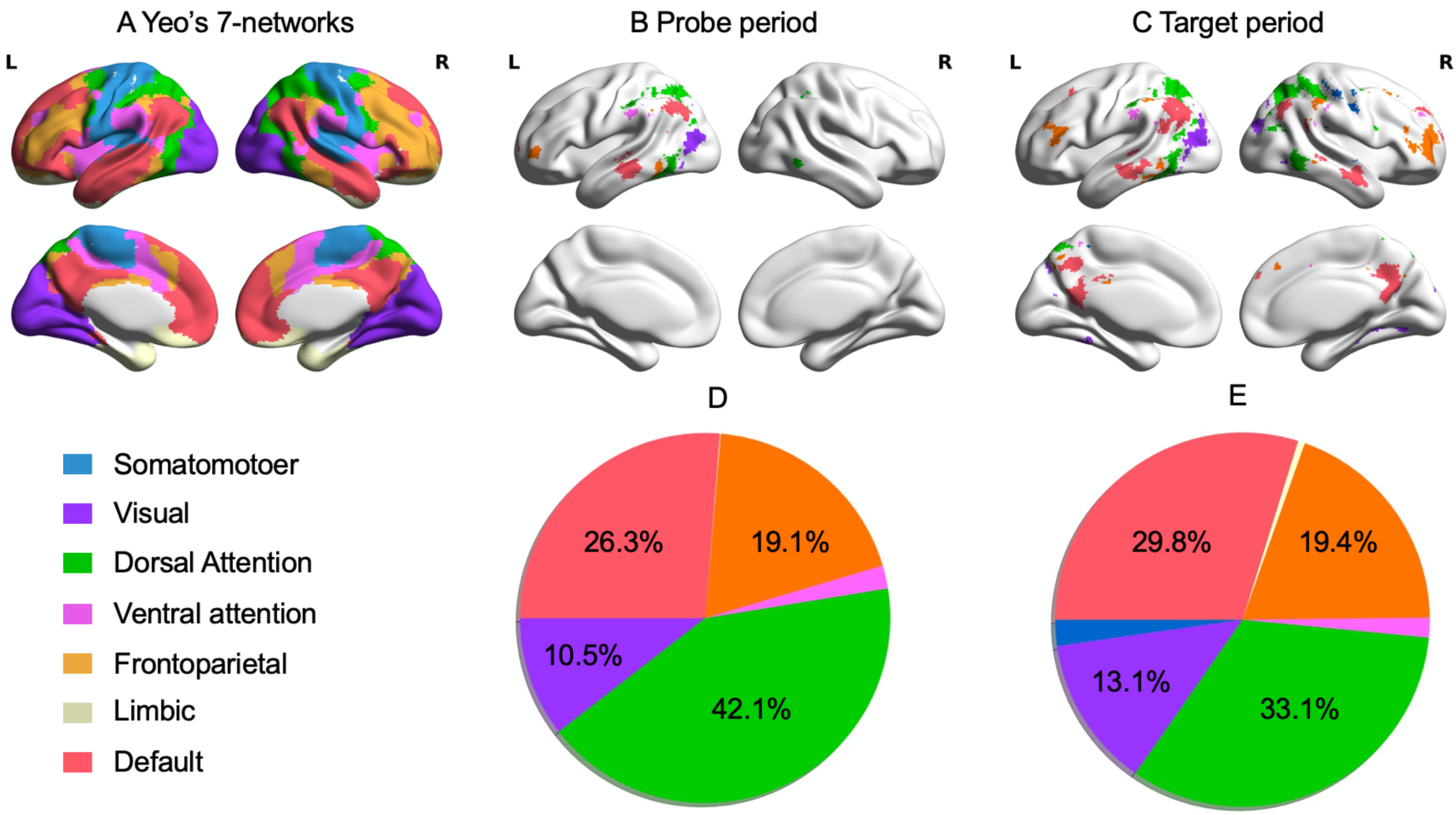
Regions representing goal information overlap with DMN and networks contributing to MDN (dorsal attention and frontoparietal networks) defined by (Yeo et al., 2011). A - The seven large scale networks identified by Yeo et al. (2011). B, C - Overlap between regions representing goal information and large scale networks during the probe and target word period, respectively. D, E - Pie chart shows the percentage of voxels within each decoding map falling within each network defined by Yeo et al. (2011) during the probe and the target word period, respectively. Values less than 3% are not shown.

To confirm this overlap of the goal decoding results with both DMN and MDN, we compared our decoding results with DMN and MDN activation maps defined for the same participants by the localizer tasks. DMN and MDN were identified using contrasts of easy and hard spatial working memory and math judgements (cf. Fedorenko et al. 2011). Consistent with previous findings, DMN regions (showing a stronger response to easy versus hard trials) included posterior cingulate cortex, medial prefrontal cortex, angular gyrus and lateral anterior temporal lobes bilaterally. In contrast, MDN regions (responding to hard versus easy trials) included inferior frontal sulcus, premotor cortex, intraparietal sulcus, and lateral occipital cortex (FWE-corrected, z = 3.1, p <.05) (Figure 5A). Regions that represented goal information overlapped with both MDN and DMN during the probe period (Figure 5B) and the target period (Figure 5C).

**Figure 5.**
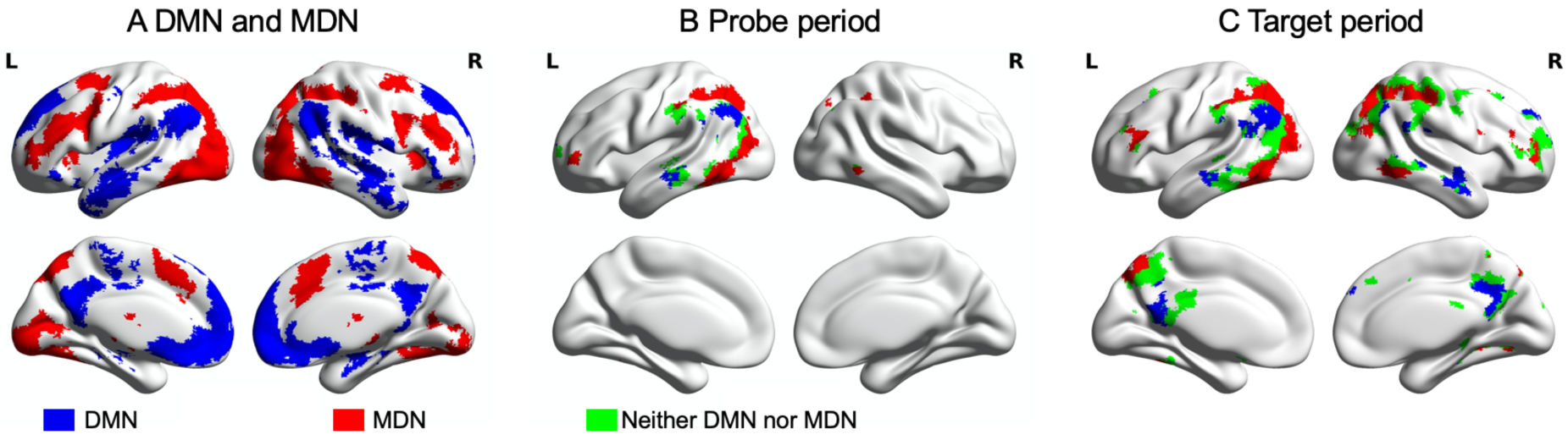
Regions representing goal information overlap with DMN and MDN defined by the localizer tasks. A - DMN and MDN defined using the localizer tasks. B, C - The overlap between regions representing goal information and DMN (blue) and MDN (red) during the probe and the target period, respectively. Regions that overlap with neither DMN nor MDN are in green.

### 3.5. Goal classification is not related to task difficulty

Activity in DMN and MDN is highly sensitive to task difficulty. Therefore, we tested for the possibility that the goal discrimination results were driven by differences in difficulty between the three goals. We calculated the correlation between the decoding accuracy in the target word period for each ROI and the absolute difference in response time between colour, shape and size trials per participant. We defined 6 ROIs by overlapping regions able to decode feature information during the target word period with corresponding ROIs defined by using a parcellation of intrinsic connectivity (Schaefer et al., 2018). These ROIs fell within the frontoparietal control network (i.e. left inferior frontal gyrus), dorsal attention network (i.e. left intraparietal sulcus, superior parietal lobe, and lateral occipital cortex) and DMN (i.e. left angular gyrus, middle temporal gyrus, and posterior cingulate cortex; Figure 6). We did not find any significant correlations between decoding accuracy and response time differences in these regions (r < 0.28, p > 0.1), suggesting that decoding accuracy was unrelated to differences in difficulty between conditions.

**Figure 6.**
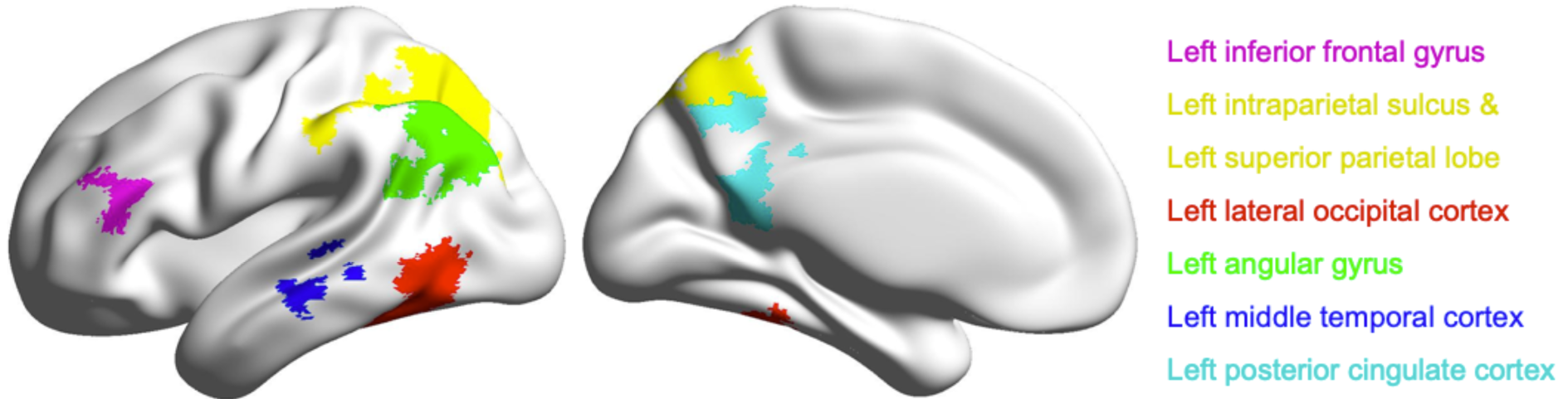
ROIs defined to examine the correlations between response difference between features and decoding accuracy.

### 3.6. Category information is represented in lateral occipital cortex

The probe words were selected from three categories – animal, tool and plant – allowing whole-brain decoding of category as well as goal information during the probe period. Probe category was decoded in left lateral occipital cortex (mean decoding accuracy = 0.362, SD = 0.027; Figure 7A). This cluster overlapped with the goal decoding results for the probe period (Figure 7B), suggesting processing in this brain region reflects both current task demands and semantic similarity.

**Figure 7.**
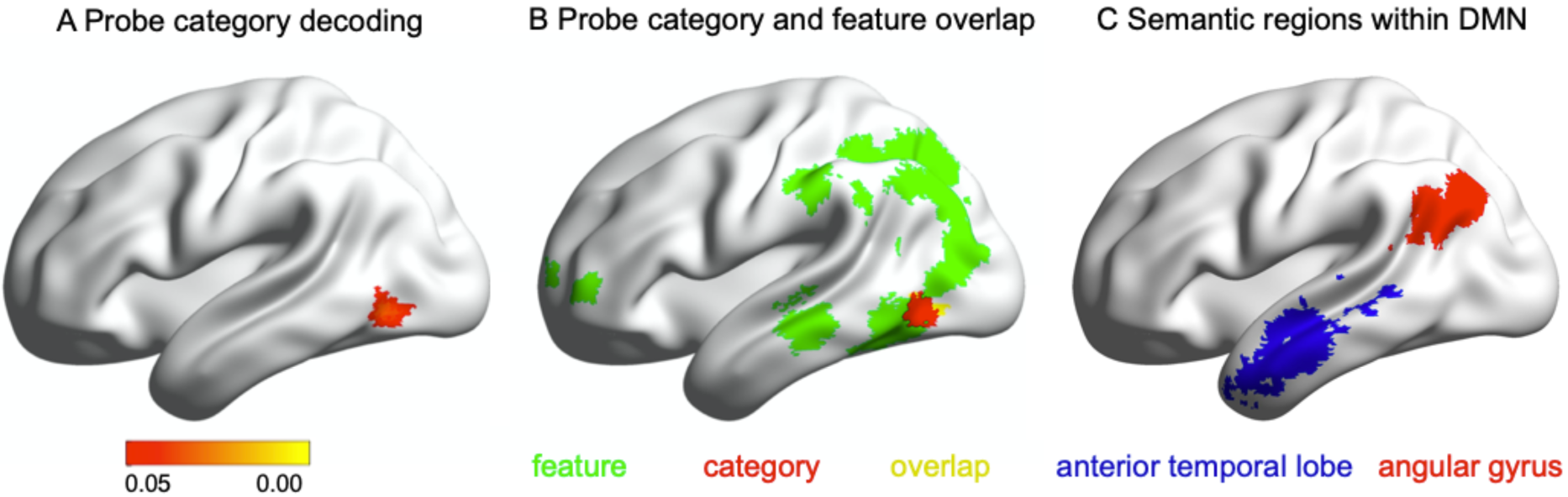
A – Region that represents category information of probe words (FWE-corrected, p < 0.05). B – Overlap of category classifier with regions that represent goal information of probe words. C – The two ROIs defined for the ROI-based classification analysis.

We reasoned that if the organisation of conceptual processing in DMN reflects long-term semantic similarity, regions of this network might represent category information, even when this not necessary for the task. The whole-brain analysis above did not identify regions of DMN that could decode semantic category. However, to guard against Type II errors, we examined whether category information could be decoded in key semantic sites within DMN by performing ROI-based classification analysis. The semantic DMN sites were defined by overlapping the DMN network defined by the localizer tasks (Figure 5A) with semantic regions defined using a meta-analytic mask for the term “semantic” from Neurosynth (Yarkoni et al., 2011). We identified two main regions for analysis in left angular gyrus and left anterior temporal lobe by overlapping these voxels with the Schaefer parcellation (Schaefer et al., 2018) (Figure 7C). We applied Bonferroni correction to account for the fact that we included two ROIs; consequently, the final threshold was p < 0.025. We found category classification was not different from the chance level (i.e. 0.33) in left angular gyrus (mean decoding accuracy = 0.346, SD = 0.013; t(27) = 1.236, p = 0.227). Classification of semantic category in the left anterior temporal lobe was trending above chance (mean decoding accuracy = 0.353, SD = 0.010; t(27) = 2.274, p = 0.031).

## 4. Discussion

This study contrasted two different accounts of DMN function. Semantic DMN sites might be expected to decode semantic similarity irrespective of task demands, consistent with their putative role as a store of heteromodal concepts. By an alternative view, DMN might show sensitivity to changing task demands, consistent with evidence that this network contains ‘echoes’ of all other networks and dynamically alters its patterns of connectivity depending on the context (Spreng et al., 2013; Vatansever et al., 2015b; Dixon et al., 2017, 2018).

We used a semantic feature matching task, which required participants to integrate conceptual knowledge with changing task goals, such that successive decisions were based on different features of the items. This task allowed us to simultaneously decode semantic category and current goal information using a whole-brain searchlight decoding approach. We replicated previous findings that regions of MDN represent current goals. We additionally demonstrated that goal information is also reflected in the multivariate response within regions of DMN, including angular gyrus and posterior cingulate cortex. Semantic category information that was not relevant to the ongoing task could be decoded in lateral occipital cortex but not regions of DMN (with the possible exception of ATL). The successful decoding of goal information alongside the absence of category decoding effects within DMN suggests that this network is sensitive to changing semantic task demands.

By examining different phases of the task, we showed that goal decoding during semantic retrieval does not reflect visual or language processes relating to the goal cue word, since these effects would have been maximised during goal cue presentation – at this time point, we found no decoding of goal beyond visual cortex. In contrast, the strong similarity between probe and target decoding results suggests that goal representations are maintained and applied to direct semantic retrieval, and that this process involves both MDN and DMN. Waskom et al. (2014) similarly found goal information could be decoded in inferior frontal sulcus within MDN when goals were applied to stimuli – however, our observation that this pattern extends to DMN is novel. Our results are also consistent with a recent study which revealed separate representational formats between goal cue and object stimulus periods (Hebart et al., 2018). While our goal decoding effects in heteromodal cortex were constrained to periods involving semantic retrieval not task preparation, in other contexts, goal representations in DMN may be critical for task preparation, for example, when participants are retrieving complex instructions from memory (Crittenden et al., 2015; Smith et al., 2018). Further research is needed to establish if DMN can decode current goals beyond situations that strongly engage memory; for example, during perceptual tasks.

Our goal classification results within regions of DMN cannot be easily explained by difficulty or spatial adjacency to MDN. There was no correlation between decoding accuracy and RT differences between goal features (e.g. colour, shape and size trials) – and DMN and MDN were both able to decode goal information, even when RT (as a proxy for difficulty) was regressed out. Moreover, although DMN and MDN often occupy adjacent brain areas (consistent with their contribution to heteromodal processing; Margulies et al., 2016), some DMN regions that are further away from MDN, such as middle temporal gyrus, were able to decode goal information in this study.

While our classification accuracies were relatively low, similar levels of performance have been commonly reported when examining goal representation across MDN (Waskom et al., 2014; Erez and Duncan, 2015; Bhandari et al., 2018). The fMRI decoding accuracy of prefrontal cortex often hovers just above chance levels, even for task features that are known to be robustly represented by the activity of prefrontal cortex neurons in nonhuman primates (Woolgar et al., 2011; Stokes et al., 2013; Bhandari et al., 2018). Whole-brain searchlight analyses typically yield even lower classification accuracies than ROI approaches (Waskom et al., 2014), explaining why many previous studies have applied an MDN mask to identify regions for decoding (Waskom et al., 2014; Erez and Duncan, 2015; Cole et al., 2016; Bracci et al., 2017; Qiao et al., 2017). This bias towards executive cortex has limited our understanding of the role of DMN in cognitive flexibility.

Our results allow us to reject accounts of the neural basis of semantic cognition that anticipate strong functional dissimilarities between DMN and MDN in all circumstances. For example, the observation that DMN tends to deactivate during demanding tasks and the deactivation is positively associated behaviour performance (Anticevic et al., 2012) has motivated the proposal that DMN is not critical for controlled cognition (Shapira-Lichter et al., 2013; Axelrod et al., 2017), or for tasks that involve attending to external inputs (even when these tasks require conceptual processing) (Chiou et al., 2020). Our findings are clearly incompatible with these views. Similarly, our results conflict with the claim that DMN regions only support relatively ‘automatic’ aspects of semantic retrieval (Vatansever et al., 2017). Although DMN regions show a greater response when strong associations are contrasted with weak associations (Davey et al., 2015; Teige et al., 2018, 2019), when meaningful inputs share a higher number of features (Wang et al., 2020) and when multiple inputs are convergent in meaning (Westerlund and Pylkkänen, 2014; Lanzoni et al., 2020), other studies find these regions also support controlled aspects of semantic cognition (Krieger-Redwood et al., 2016; Murphy et al., 2018), in line with our observation that DMN regions can decode current goals.

Other accounts of DMN have emphasised functional subsystems, identified through patterns of intrinsic connectivity and functional recruitment (Andrews-Hanna et al., 2010). A recent study found that “core” DMN regions (e.g. angular gyrus and posterior cingulate cortex) show more task-related deactivation and selective recruitment for ‘internal cognition’, compared with lateral temporal DMN regions implicated in semantic processing (Chiou et al., 2020). By this view, the task-negative and controlled memory accounts of DMN can be reconciled by attributing these patterns of functional recruitment to different DMN subsystems. Our findings are not compatible with this account, since our goal decoding results extended over both DMN subsystems: both core regions (such as posterior cingulate cortex) and lateral temporal DMN regions could classify task goals during semantic retrieval. Our findings are more consistent with recent findings that DMN and MDN have functional similarities, despite their well-documented activation differences. For example, these brain regions can have similar representational formats (González-García et al., 2018) and they occupy adjacent positions on the principal gradient of connectivity, which explains the most variance in a decomposition of resting-state fMRI (Margulies et al., 2016; Mckeown et al., 2020)

An adequate account of the role of DMN in semantic cognition needs to explain the stronger activation typically seen in this network when the meanings of inputs are well-aligned with recent experience or long-term memory, as well as the sensitivity of this network to changing task goals. One possibility is provided by views that envisage DMN regions as “integrative hubs” (Braga et al., 2013), drawing together inputs from highly diverse networks, including unimodal regions relevant to the varied features of concrete concepts (e.g., colour, shape, size) (Margulies et al., 2016; Lanzoni et al., 2020). In this framework, each feature matching task requires a specific pattern of interactive processing between DMN hubs and specific unimodal regions relevant for the task. The distinct multivariate responses that we uncovered within DMN for each task goal might have corresponded to these different task states. In this way, our ability to decode changing task goals in DMN might reflect the location of this network at the top of a cortical hierarchy extending from unimodal to heteromodal cortex (Margulies et al., 2016), explaining the engagement of this network across a wide variety of tasks. At the same time, cognitive states involving convergent features across multiple unimodal regions might be expected to elicit strong univariate responses within the same DMN regions (Lanzoni et al., 2020; Wang et al., 2020), since this joining together of inputs would drive strong pattern completion properties across the cortex.

Although semantically-relevant DMN regions were unable to decode category when this was not relevant to the ongoing task, lateral occipital cortex could decode both category and goal. Similar effects have been reported previously, reflecting strong visual similarities amongst category members – for example, animals tend to be similar shapes (Bracci et al., 2017; Bugatus et al., 2017). Consequently, it has been proposed that visual responses in lateral occipital cortex are largely context and task invariant (Bracci et al., 2017; Bugatus et al., 2017; Xu, 2018a, 2018b). However, lateral occipital cortex is also commonly-activated across multiple control-demanding tasks, and by this definition, forms part of the MDN (Waskom et al., 2014; Gonzalez Alam et al., 2018; Hebart et al., 2018). Behavioural goals directly impact object representations in this region (Harel et al., 2014). Consequently, our results support the view that lateral occipital cortex represents a mixture of task and object information (Hebart et al., 2018). Further research is needed to establish the circumstances in which DMN regions might similarly show decoding of both content and goal, rather than goal alone.

## Acknowledgement

This study was supported by the European Research Council (Project ID: 771863 - FLEXSEM and Project ID: 646927 - WANDERINGMINDS). We are grateful to Evelina Fedorenko for providing us with the localizer tasks and to Evgeny Gluzman for piloting a preliminary version of the task. We thank Charlotte Murphy, Jiefeng Jiang, Deniz Vatansever and Theodoros Karapanagiotidis for suggestions about data analysis.

